# Buprenorphine Induces Human Fetal Membrane Sterile Inflammation

**DOI:** 10.1101/2024.11.22.624850

**Authors:** Tatyana Lynn, Megan E Kelleher, Hanah M Georges, Elle M McCauley, Ryan W Logan, Kimberly A Yonkers, Vikki M Abrahams

## Abstract

Opioid-use disorder (OUD) during pregnancy has increased in the United States to critical levels and is a leading cause of maternal morbidity and mortality. Untreated OUD is associated with pregnancy complications in particular, preterm birth. Medications for OUD, such as buprenorphine, are recommended with the added benefit that treatment during pregnancy increases treatment post-partum. However, the rate of preterm birth in individuals using illicit opioids or being treated with opioid agonist therapeutics is double that of the general population. Since inflammation in the placenta and the associated fetal membranes (FM) is a common underlying cause of preterm birth, we sought to determine if the opioid, buprenorphine, induces sterile inflammation in human FMs and to examine the mechanisms involved. Using an established *in vitro* human FM explant system, we report that buprenorphine significantly increased FM secretion of the inflammatory cytokine IL-6; the neutrophilic chemokine IL-8; and the inflammasome-mediated cytokine IL-1β, mirroring the inflammatory profile commonly seen at the maternal-fetal interface in preterm birth. Other factors that were elevated in FMs exposed to buprenorphine included the mediators of membrane weakening, prostaglandin E2 (PGE2), and matrix metalloproteinases, MMP1 and MMP9. Furthermore, this sterile inflammatory and weakening FM response induced by buprenorphine was mediated in part by innate immune Toll-like receptor 4 (TLR4), the NLRP3 inflammasome, the μ-opioid receptor, and downstream NFκB and ERK/JNK/MAPK signaling. This may provide the mechanistic link between opioid use in pregnancy and the elevated risk for preterm birth. Since there are adverse consequences of not treating OUD, our findings may help identify ways to mitigate the impact opioids have on pregnancy outcomes while allowing the continuation of maintenance therapy.

## Introduction

Opioid-use disorder (OUD) during pregnancy is a leading cause of maternal morbidity and mortality [1–3]. Untreated OUD is associated with pregnancy complications including preterm birth, fetal growth restriction (FGR) and fetal demise [2]. Standard care for OUD is maintenance therapy with buprenorphine or methadone to supply a steady concentration of opioid throughout treatment in order to avoid the risk of relapse, additional disease from needle sharing, and unregulated drug use [4, 5]. An added benefit of treatment during pregnancy is increased treatment post-partum [1, 6, 7]. Although methadone and buprenorphine are used to treat OUD, patients taking these opioids still have an increased risk of preterm birth, demonstrating that any opioid use can adversely affect pregnancy outcomes. In one study, the preterm birth rate for individuals taking buprenorphine was as high as 22.7% [2] compared to 10.4% in the general population [8].

Preterm birth is a major contributor to neonatal morbidity and mortality, and is often associated with the presence of chorioamnionitis - inflammation of the fetal membranes (FM) [9]. While infection-associated inflammation is a known underlying cause of preterm birth [10, 11], during chorioamnionitis and preterm birth the FM can also produce pro-inflammatory factors by sensing and responding to non-infectious triggers [12, 13]. In both cases, innate immune receptors such Toll-like receptors (TLRs) or inflammasome family members, can induce FM inflammation and preterm birth [12, 14–16]. Inflammation-driven production of mediators of membrane weakening such as prostaglandins (PGE) and matrix metalloproteinases (MMPs) can deteriorate the integrity of the membrane leading to rupture [17, 18].

Currently, there is little known about the effects of opioid exposure on human FMs or the mechanisms involved. However, the opioids, buprenorphine and methadone, have been found to induce inflammation through TLR4/MD2 and TLR2 activation in brain cells [19]. In this current study we sought to determine the direct impact of buprenorphine on human FM inflammation and mediators of membrane weakening and examine the mechanism involved. Herein, we report that the opioid, buprenorphine, induces human FM sterile inflammation and mediators of membrane weakening through activation of TLR4/MD2, the NLRP3 inflammasome, the μ-opioid receptor, and downstream NFκB and ERK/JNK/MAPK signaling. This may provide the mechanistic link between opioid use in pregnancy and the elevated risk for preterm birth. Since there are adverse consequences of not treating OUD, our findings may help identify ways to mitigate the impact opioids have on pregnancy outcomes while allowing the continuation of maintenance therapy.

## Materials and Methods

### Patient Samples

Human fetal membrane (FM) tissue collection was approved by the Yale University’s Human Research Protection Program (IRB# 0607001625) and was facilitated by the Yale University Reproductive Sciences (YURS) Biobank (IRB# 1309012696) with patient consent. Human fetal membranes were collected from uncomplicated term pregnancies (38-41 weeks’ gestation), delivered by scheduled caesarean section, in the absence of labor, known infection or known medication/opioid use, and were transferred to the laboratory immediately after delivery.

### Treatment of FMs

An established FM explant system was used [15]. After washing FMs with PBS containing 1% penicillin (100 U/ml) and streptomycin (100μg/ml) (Life Technologies, Grand Island, NY), 6mm explants were placed in 0.4μm cell culture inserts. The chorion and amnion were kept intact. Inserts were placed in 24 well plates containing 1ml (500μl in the upper and lower chambers) of Dulbecco modified Eagle medium (DMEM; Life Technologies) supplemented with 10% FBS for overnight incubation. The following day, DMEM media was removed and replaced with serum-free OptiMeM (Life Technologies). FMs were treated with either no treatment media (NT) or buprenorphine (Covetrus, Dublin, Ohio). For some experiments, FMs were treated with NT or buprenorphine in the presence or absence of inhibitors to: TLR4 (LPS-RS, 10μg/ml) (Invivogen, San Diego, CA); TLR2 (TL2-C29, 50μM) (Invivogen, San Diego, CA); p38 MAPK (SB203580, 1μM); p65 NFκB (BAY11-7085, 10μM) (Invivogen, San Diego, CA); ERK (SCH77298, 10nM) (Selleck Chemicals, Houston, TX); JNK (SP600125, 25μM) (Millipore Sigma, Burlington, VT), and NLRP3 (MCC950, 10μM) (Invivogen, San Diego, CA). FMs were also treated with NT or buprenorphine in the absence or presence of (-) Naloxone (20μg/ml and 40μg/ml) (Thomas Scientific, Swedesboro, NJ). Inhibitors and (-) naloxone were added to FM culture system 1hr prior to buprenorphine treatment. After 24hrs or 48hrs, cell-free supernatants and FM tissues were collected. Tissues were snap frozen. Supernatants were centrifuged to remove any cellular debris. Both tissues and supernatants were stored at -80°C. All FM experiments were performed in triplicate.

### Analysis of FM supernatants for inflammatory markers

FM supernatants were measured by ELISA for IL-6, IL-8, IL-1β, G-CSF, PGE2, MMP1, MMP2, MMP9, CCL7/MCP3, CSF2/GM-CSF, CXCL6, CXCL5/RANTES, CCL3/MIP-1α, TNF-α, and CCL2/MCP-1 (R&D Systems, Minneapolis, MN). All assays were run in duplicate.

### Western Blot Analysis

FM tissues were homogenized for protein in lysis buffer containing phosphatase inhibitors (Cell Signaling) and concentrations measured using the Pierce BCA protein assay (ThermoFisher Scientific, Waltham, MA). Membranes were probed for the following primary antibodies all from Cell Signaling Technology (Danvers, MA): phosphorylated (p)-p65 NFκB (#3033; 1:1000); total (t)-p65 NFκB (#8242; 1:1000); p-p38 MAPK (#9211;1:500); t-p38 MAPK (#9212; 1:1000); p-ERK (#9101; 1:1000); t-ERK (#4695; 1:1000); p-JNK (#9251; 1:1000); t-JNK (#; 1:1000); Lamin B1 (#1343; 1:1000); p16 (#92803; 1:1000). β-Actin (A2066; 1:10000) from Sigma Aldrich was used as a loading control for Lamin B1 and p16. Western blot images were captured using an Amersham Imager 680 (General Electric, Boston, MA). Image Studio Lite (Li-Cor Biosciences, Lincoln, NE) was used to perform semiquantitative densitometry. Fold changes were determined by normalizing phosphorylated protein levels against the paired total protein levels. For LaminB1 and p16, proteins levels were normalized against β-Actin.

### Bulk RNA-Seq Analysis

RNA was extracted using TRIzol (ThermoFisher Scientific; Waltham, MA) according to manufacturer’s protocol. Samples were submitted to the Yale Center for Genome Analysis (YCGA) for RNA sequencing using the following methods: Total RNA quality was determined by estimating the A260/A280 and A260/A230 ratios by Nanodrop Spectrophotometer (ThermoFisher Scientific; Waltham, MA). RNA integrity was determined by running an Agilent Bioanalyzer gel, which measures the ratio of the ribosomal peaks.

#### RNA Seq Library Prep

Using the Kapa RNA HyperPrep Kit with RiboErase (KR1351), rRNA was depleted starting from a normalized input of total RNA by hybridization of rRNA to complementary DNA oligonucleotides, followed by treatment with RNase H and DNase to remove rRNA duplexed to DNA. Samples were then fragmented using heat and magnesium. First strand synthesis was performed using random priming. Second strand synthesis incorporated dUTPs into the second strand cDNA. Adapters were then ligated and the library amplified. Strands marked with dUTPs were not amplified allowing for strand-specific sequencing. Indexed libraries that met appropriate cut-offs for both quantity and quality were quantified by RT-qPCR using a commercially available kit (KAPA Biosystems; Roche Basel, Switzerland) and insert size distribution determined with the LabChip GX or Agilent Bioanalyzer.

#### Flow Cell Preparation and Sequencing

Sample concentrations were normalized to 2.0 nM and loaded onto an Illumina NovaSeq flow cell at a concentration that yields 35 million passing filter clusters per sample. Samples were sequenced using 100bp paired-end sequencing on an Illumina NovaSeq (San Diego, CA) according to Illumina protocols. The 10bp unique dual index was read during additional sequencing reads that automatically follow the completion of read 1. Data generated during sequencing runs were simultaneously transferred to the YCGA high-performance computing cluster. A positive control (prepared bacteriophage Phi X library) provided by Illumina was spiked into every lane at a concentration of 0.3% to monitor sequencing quality in real time.

#### Data Analysis and Storage

Signal intensities were converted to individual base calls during a run using the system’s Real Time Analysis (RTA) software. Base calls were transferred from the machine’s dedicated personal computer to the Yale High Performance Computing cluster via a 1 Gigabit network mount for downstream analysis. Primary analysis - sample de-multiplexing and alignment to the human genome - was performed using Illumina’s CASAVA 1.8.2 software suite. Ingenuity Pathways Analysis (IPA; Qiagen Hilden, Germany) was utilized and pathways with the highest statistical significance were selected for evaluation.

### Statistical Analysis

Each *in vitro* FM experiment utilized a single patient FM. FM explants were prepared and treated in triplicate, analyzed in duplicate, and data averaged. Data are represented as a mean ± standard error of mean (SEM) from pooled experiments. Statistical significance (*p*<0.05) was determined by performing, for normalized data one-way ANOVA or a paired T-test, or if not normally distributed, a non-parametric multiple comparison test or the Wilcoxon matched-pairs signed rank test using Prism Software (Graphpad, Inc, La Jolla, CA).

## Results

### Buprenorphine induces human FM sterile inflammation

To determine the effects of buprenorphine on human FM sterile inflammation, the secretory levels of three pro-inflammatory factors that are associated with preterm birth were measured: the inflammatory cytokine IL-6; the neutrophilic chemokine IL-8; and the inflammasome-mediated cytokine IL-1β [20, 21]. As shown in Figure 1A, buprenorphine at a dose of 40μg/ml significantly increased human FM explant secretion of IL-6 by 4.3±1.2-fold, IL-8 by 2.4±0.1-fold, and IL-1β by 4.6±0.4-fold when compared to the media (0μg/ml) control. The lower doses of buprenorphine that were tested (10μg/ml and 20μg/ml) had no significant effect on the secretion of these factors by FM explants (Figure 1A). Based on this, we continued our studies using buprenorphine at 40μg/ml which is an *in vitro* dose similar to that used by others [19, 22, 23]. Since we found in a previous study that sterile FM inflammation in response to an antidepressant also elevated the secretion of G-CSF [13], we analyzed the production of this factor as well as the markers of membrane weakening, PGE2, MMP1, MMP2 and MMP9 [13]. As shown in Figure 1B, buprenorphine at a dose of 40μg/ml significantly increased human FM explant secretion of G-CSF by 5.0±0.1-fold, PGE2 by 67.6±35.0-fold, MMP1 by 2.86±1.1-fold, and MMP9 by 1.6±0.2-fold when compared to the no treatment (NT) control, while MMP2 levels were not significantly changed.

**Figure 1.**
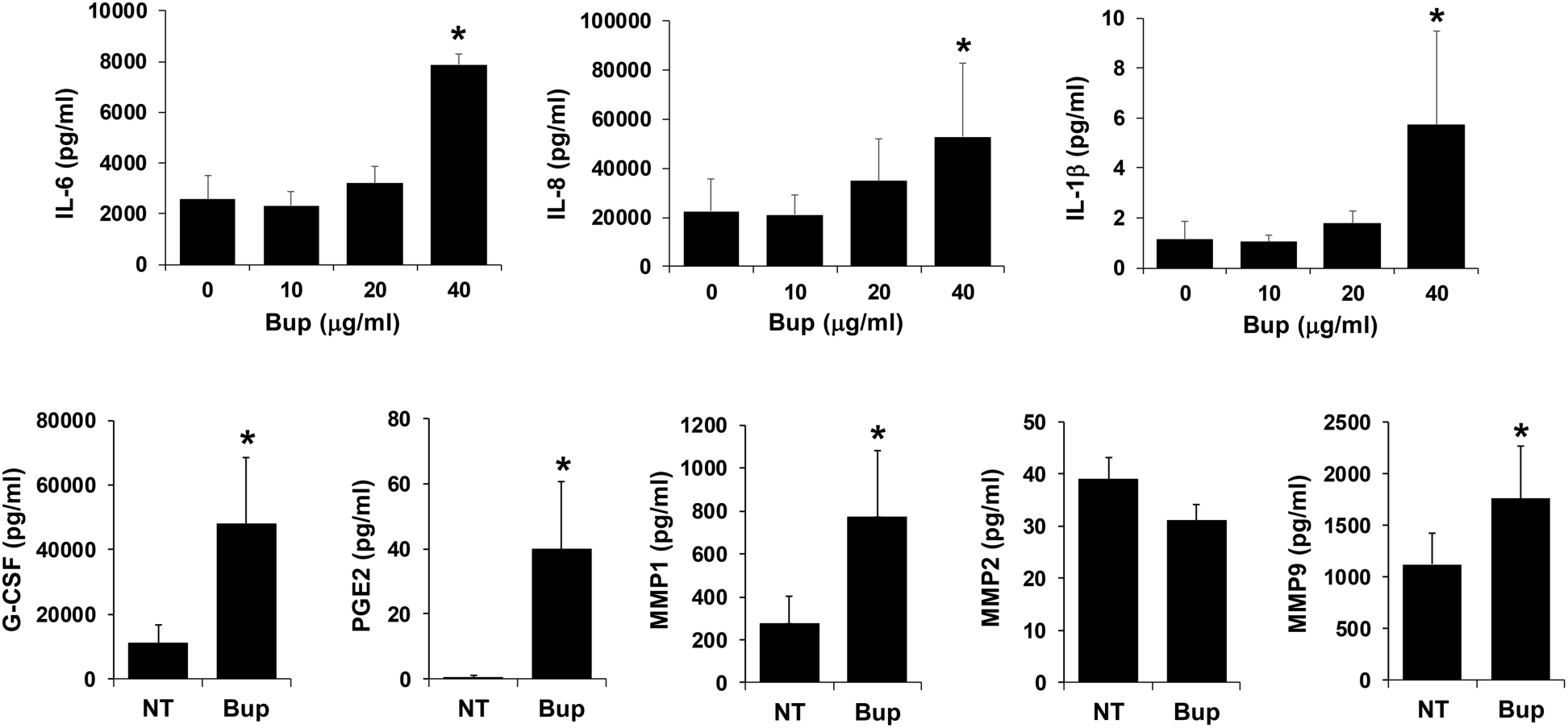
Buprenorphine induces human FM sterile inflammation. Human FM explants from 4-10 patients were treated with (A) buprenorphine at 0, 10, 20 or 40μg/ml; or (B) no treatment (NT) or buprenorphine (Bup) at 40μg/ml. After 48hrs, cell-free supernatants were collected and measured by ELISA for (A) IL-6, IL-8, and IL-1β; and (B) G-CSF, PGE2, MMP1, MMP2 and MMP9. \**p*<0.05 relative to NT control or buprenorphine at 0μg/ml.

### Buprenorphine-induced human FM inflammation is partially TLR4 dependent, but independent of TLR2 activation

Given that buprenorphine has been shown to active TLR4/MD2 and TLR2 to induce inflammation in brain cells [19, 24], we sought to determine if these innate immune receptors were involved in the FM inflammatory response to this opioid. The TLR4/MD2 antagonist, LPS-RS, significantly inhibited buprenorphine-induced FM secretion of IL-6 by 19.0±8.5% and G-CSF by 26.0±10.1%, without affecting the ability of buprenorphine to elevate FM secretion of IL-8, IL-1β, G-CSF, PGE2, MMP1 and MMP9 (Figure 2A). The TLR2 inhibitor, TL2-C29, had no effect on the ability of buprenorphine to induce a FM inflammatory response (Figure 2B).

**Figure 2.**
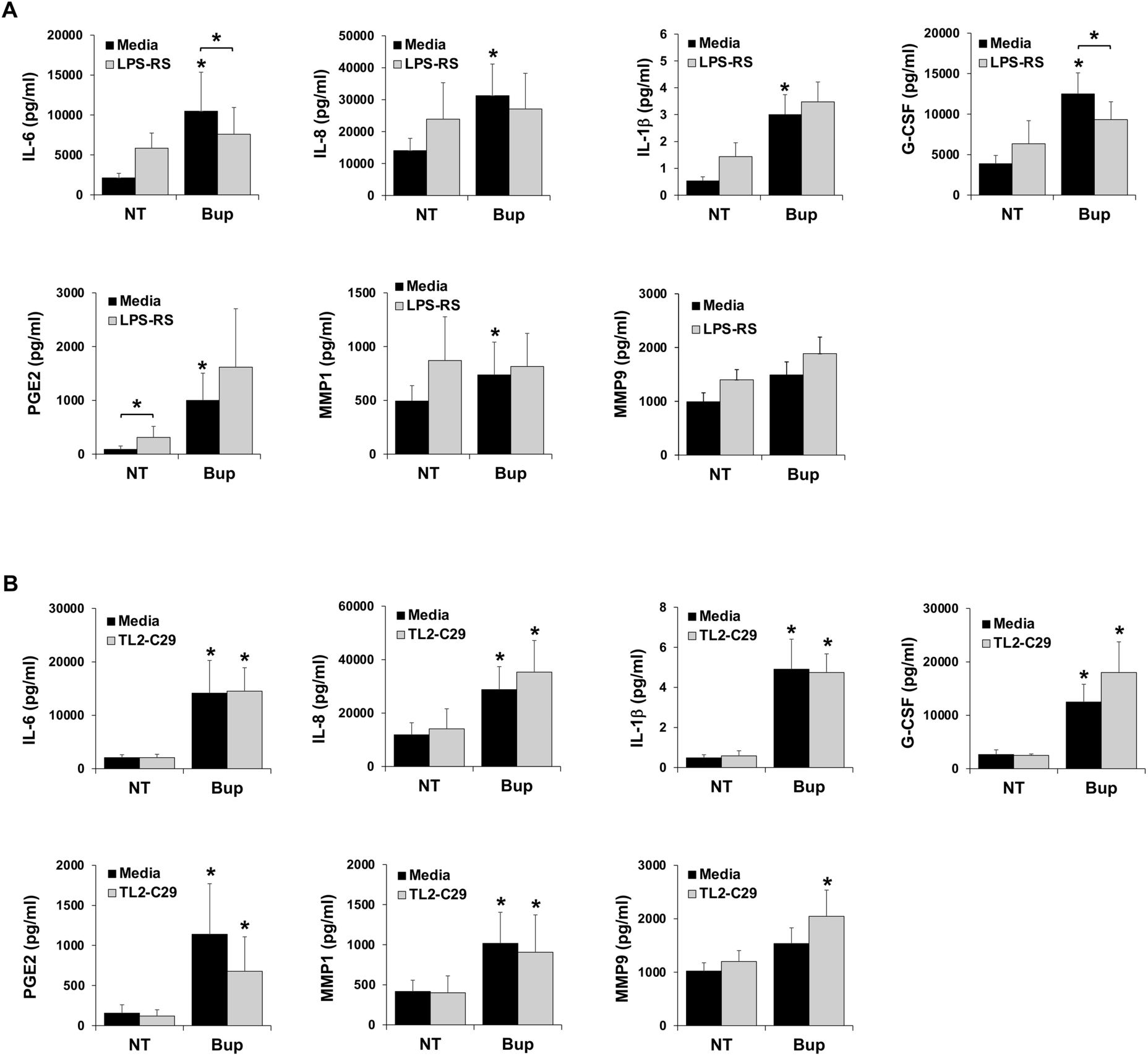
Buprenorphine induced human FM inflammation is partially TLR4 dependent but independent of TLR2 activation. Human FM explants from 7-10 patients were treated with no treatment (NT) or buprenorphine (Bup, 40μg/ml) in the presence of media or (A) the TLR4 inhibitor LPS-RS (10μg/ml) or (B) the TLR2 inhibitor, TL2-C29 (50μM). After 48hrs, cell-free supernatants were collected and measured by ELISA for IL-6, IL-8, IL-1β, G-CSF, PGE2, MMP1, and MMP9. \**p*<0.05 relative to NT/media or NT/inhibitor control or as otherwise indicated.

### Buprenorphine induced human FM inflammation is in part mediated by the μ-opioid receptor

Buprenorphine is a partial mu (μ)-opioid receptor agonist. The opioid receptors κ, μ and δ are expressed by trophoblasts [25] which constitute the majority of FM chorionic cells [26, 27]. There is evidence that opioid receptors can mediate inflammatory signaling [25]. Further, the opioid receptor antagonist, (-) naloxone [28], inhibits not only the opioid receptor, but also TLR4/MD2-mediated inflammation [29, 30]. As shown in Figure 3, (-) naloxone at 20μg/ml significantly reduced buprenorphine-induced FM secretion of IL-6 by 16.5±6.4% and IL-8 by 22.3±11.5%, while (-) naloxone at 40μg/ml significantly reduced buprenorphine-induced FM secretion of IL-8 by 16.8±7.5%, and PGE2 by 34.1±15.9%. Naloxone had no significant effect on the ability of buprenorphine to elevate FM secretion of IL-1β, G-CSF, MMP1 or MMP9 (Figure 3).

**Figure 3.**
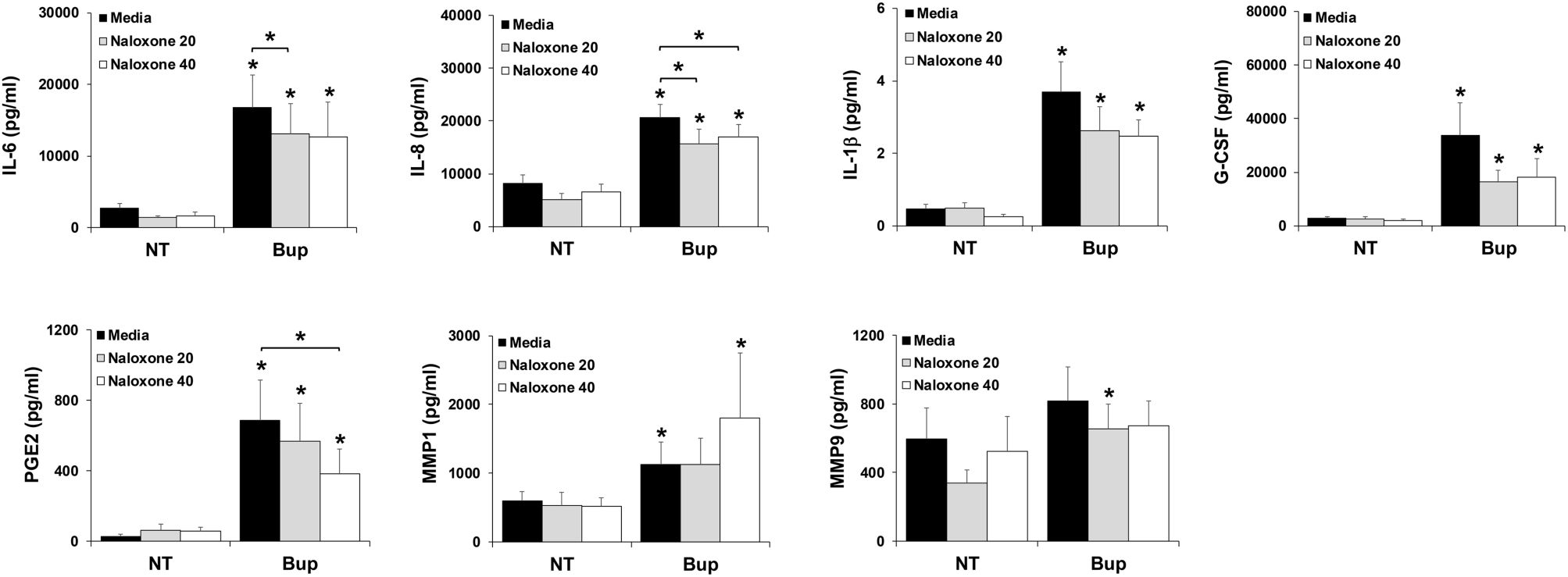
Buprenorphine induced human FM inflammation is in part mediated by the μ-opioid receptor. Human FM explants from 13 patients were treated with no treatment (NT) or buprenorphine (Bup, 40μg/ml) in the presence of media or the opioid receptor antagonist (-) naloxone at either 20μg/ml or 40μg/ml. After 48hrs, cell-free supernatants were collected and measured by ELISA for IL-6, IL-8, IL-1β, G-CSF, PGE2, MMP1, and MMP9. \**p*<0.05 relative to NT/media or NT/naloxone controls, or as otherwise indicated.

### Buprenorphine induces human FM IL-1β production through activation of the NLRP3 inflammasome

Activation of the NLRP3 inflammasome plays an important role in FM production of IL-1β [15]. As shown in Figure 4, the presence of the NLRP3 inhibitor, MCC950, significantly reduced buprenorphine-induced FM secretion of IL-1β by 44.6±6.0%.

**Figure 4.**
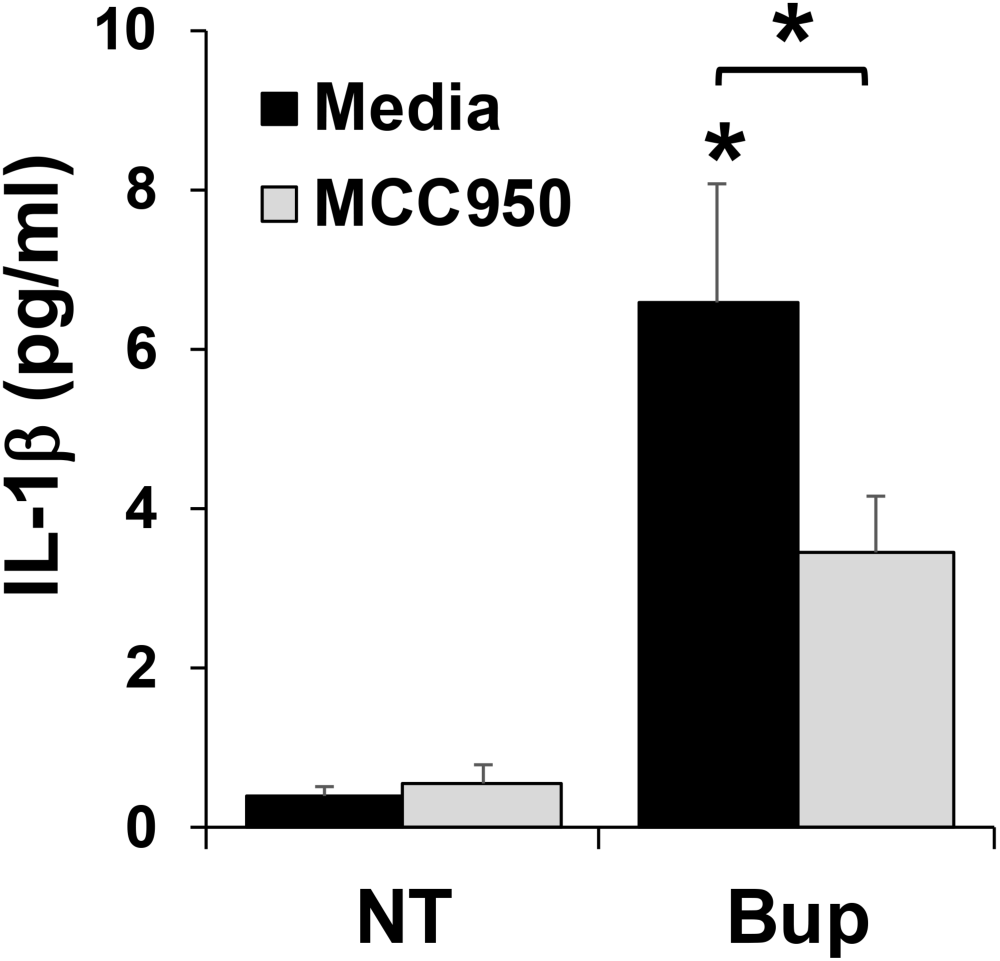
Buprenorphine induces human FM IL-1β production through activation of the NLRP3 inflammasome. Human FM explants from 6 patients were treated with no treatment (NT) or buprenorphine (Bup, 40μg/ml) in presence of media or the NLRP3 inhibitor, MCC950. After 48hrs, cell-free supernatants were collected and measured by ELISA IL-1β. \**p*<0.05 relative to NT/media or NT/inhibitor controls, or as otherwise indicated.

### Buprenorphine induced human FM inflammation is partially mediated by the p65 NFκB, p38 MAPK, ERK, and JNK signaling pathways

Having established that buprenorphine induces FM sterile inflammation and mediators of membrane weakening in part through activation of TLR4, NLRP3 and the μ-opioid receptor, we next sought to investigate the downstream signaling pathways involved. Western blot analysis showed that after 24hrs, buprenorphine elevated human FM tissue expression of phosphorylated p65 NFκB (4.4±1.4-fold) and JNK (2.3±0.7-fold), and after 48hrs buprenorphine elevated human FM tissue expression of phosphorylated p38 MAPK (1.7±0.2-fold) and ERK (3.4±1.0-fold) (Figure 5). Since FM p38 MAPK activation is associated with accelerated FM senescence [31], we evaluated the expression levels of the markers Lamin B1 and p16. As shown in Figure 5, exposure of FMs to buprenorphine after 48hrs had no effect on the expression levels of Lamin B1 or p16. This was also the case after 24hrs (data not shown).

**Figure 5.**
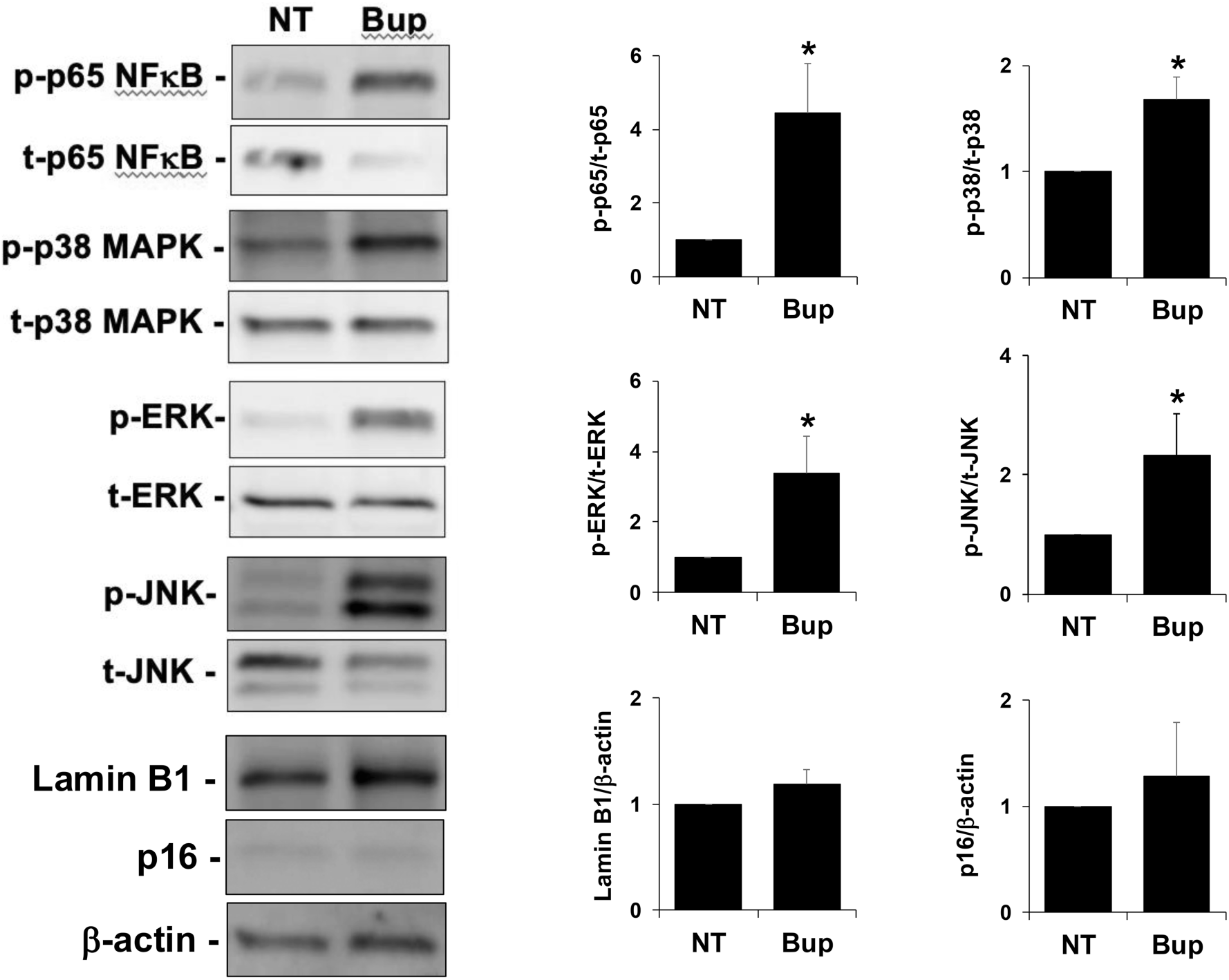
Buprenorphine activates human FM p65 NFκB, p38 MAPK, ERK and JNK signaling pathways, but does not induce senescence. Human FM explants were treated with no treatment (NT) or buprenorphine (Bup, 40μg/ml) for 24hrs (n=9) or 48hrs (n=3-6), after which tissues were homogenized for protein and Western Blot Analysis performed. Images are from representative blots. p-p65/t-p65 and p-JNK/t-JNK expression was measured at 24hrs; all other proteins were measured at 48hrs. Bar charts show quantification of protein expression as determined by densitometry (\**p*<0.05; FC= fold change).

When the p65 NFκB pathway was blocked using the inhibitor, BAY11-7082, FM secretion of IL-6, IL-8, and G-CSF in response to buprenorphine was reduced by 62.3±8.6%, 38.7±12.4%, and 40.9±13.7%, respectively, while the levels of PGE2 and MMP1 were unchanged (Figure 6A). Interestingly, inhibition of p65 NFκB significantly augmented the secretion of IL-1β by FMs exposed to buprenorphine (Figure 6A). When the p38 MAPK pathway was blocked using the inhibitor, SB203580, FM secretion of IL-6 and MMP1 in response to buprenorphine was reduced by 72.8±4.5% and 41.6±7.7%, respectively (Figure 6B). FM secretion of IL-8, IL-1β, G-CSF and PGE2 were not significantly changed by the presence of the P38 MAPK inhibitor (Figure 6B). The presence of the ERK inhibitor, SCH1772984, significantly reduced the ability of buprenorphine to upregulate FM secretion of IL-6 (63.9±6.4%), IL-8 (50.1±11.6%), G-CSF (66.4±9.1%), and MMP1 (88.3±6.5%), without affecting the secretion of IL-1β, and PGE2 (Figure 6C). The only secreted FM factor tested to be downregulated by the JNK inhibitor in the presence of buprenorphine was PGE2 which was reduced by 73.8±14.3% (Figure 6D).

**Figure 6.**
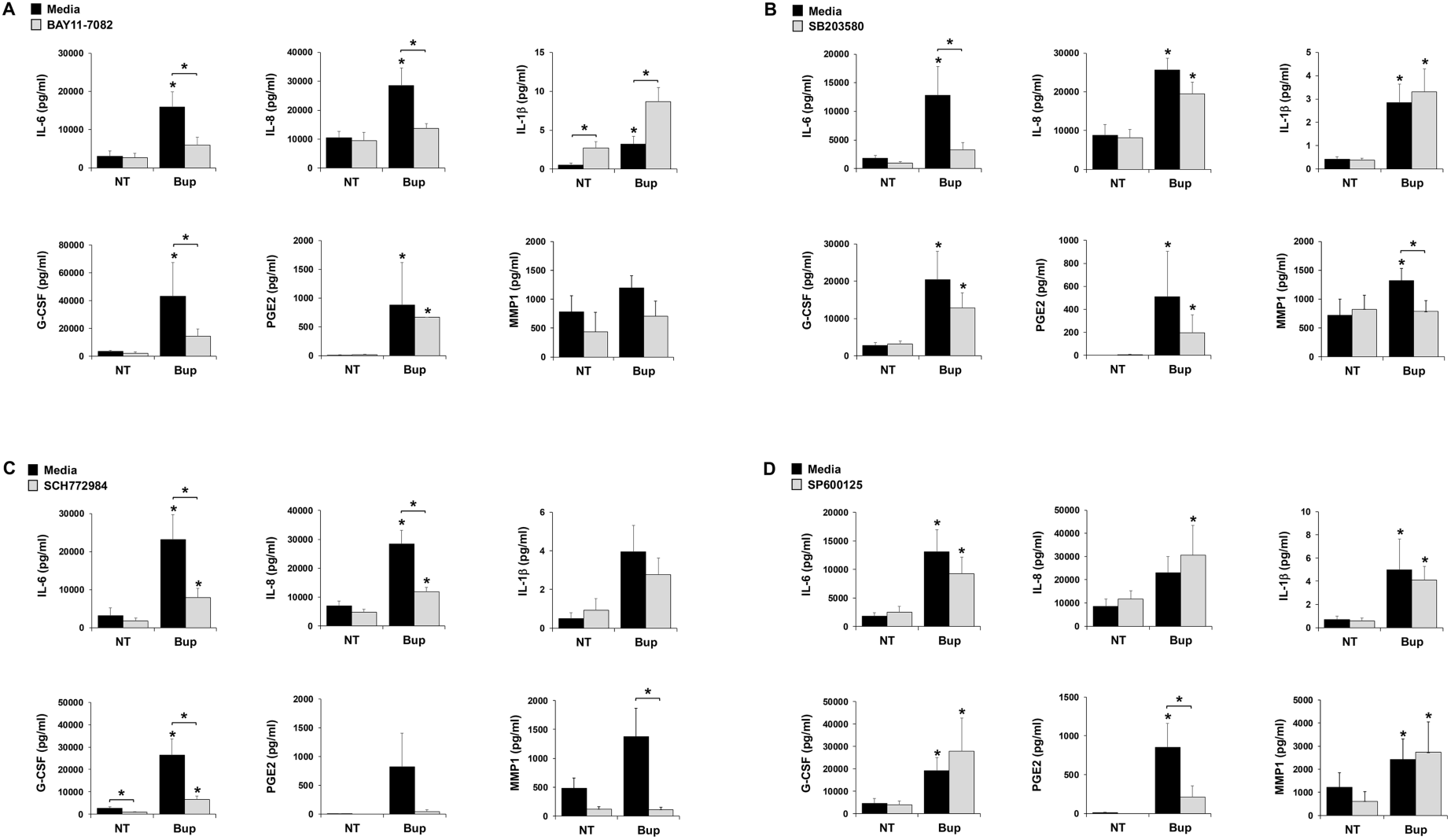
Buprenorphine induced human FM inflammation is partially mediated by the p65 NFκB, p38 MAPK, and ERK signaling pathways. Human FM explants from 6-7 patients were treated with no treatment (NT) or buprenorphine (Bup, 40μg/ml) in the presence of media or (A) the p65 NFκB inhibitor, BAY11-7085; (B) the p38 MAPK inhibitor, SB203580; (C) the ERK inhibitor, SCH77298; or (D) the JNK inhibitor, SP600125. After 48hrs, cell-free supernatants were collected and measured by ELISA for IL-6, IL-8, IL-1β, G-CSF, PGE2, MMP1, and MMP9. \**p*<0.05 relative to NT/media or NT/inhibitor controls, or as otherwise indicated.

### Validation of RNA-seq and Ingenuity Pathways Analysis

When RNA-seq was performed on FM tissues that were either not treated (NT) or exposed to buprenorphine, a significant pathway highlighted from Ingenuity Pathways Analysis was “Cytokine Storm” (Figure 7A & Supplemental Figure 1). Within this pathway, IL6, CXCL8 (IL8), and IL1B were in the top 20 genes that were significantly unregulated in response to buprenorphine, thus validating our original ELISA data (Figure 7B). From this list of upregulated genes, we selected 7 more to validate based on their high level of fold change (CCL7/MCP3, CSF2/GM-CSF, CXCL6), relevance to preterm birth (TNF), and relevance to chemotaxis and inflammation (CXCL5/RANTES, CCL3/MIP1A, CCL2/MCP1) (Figure 7B). As shown in Figure 7C, validation of these 7 factors by ELISA showed that treatment of FMs with buprenorphine significantly elevated the secretion of CSF2/GM-CSF, CXCL5/RANTES, CCL3/MIP1A, TNF-α, and CCL2/MCP-1. The secretion of only 2 factors, CCL7/MCP3 and CXCL6, were not significantly altered by buprenorphine treatment of FMs (Figure 7C).

**Figure 7.**
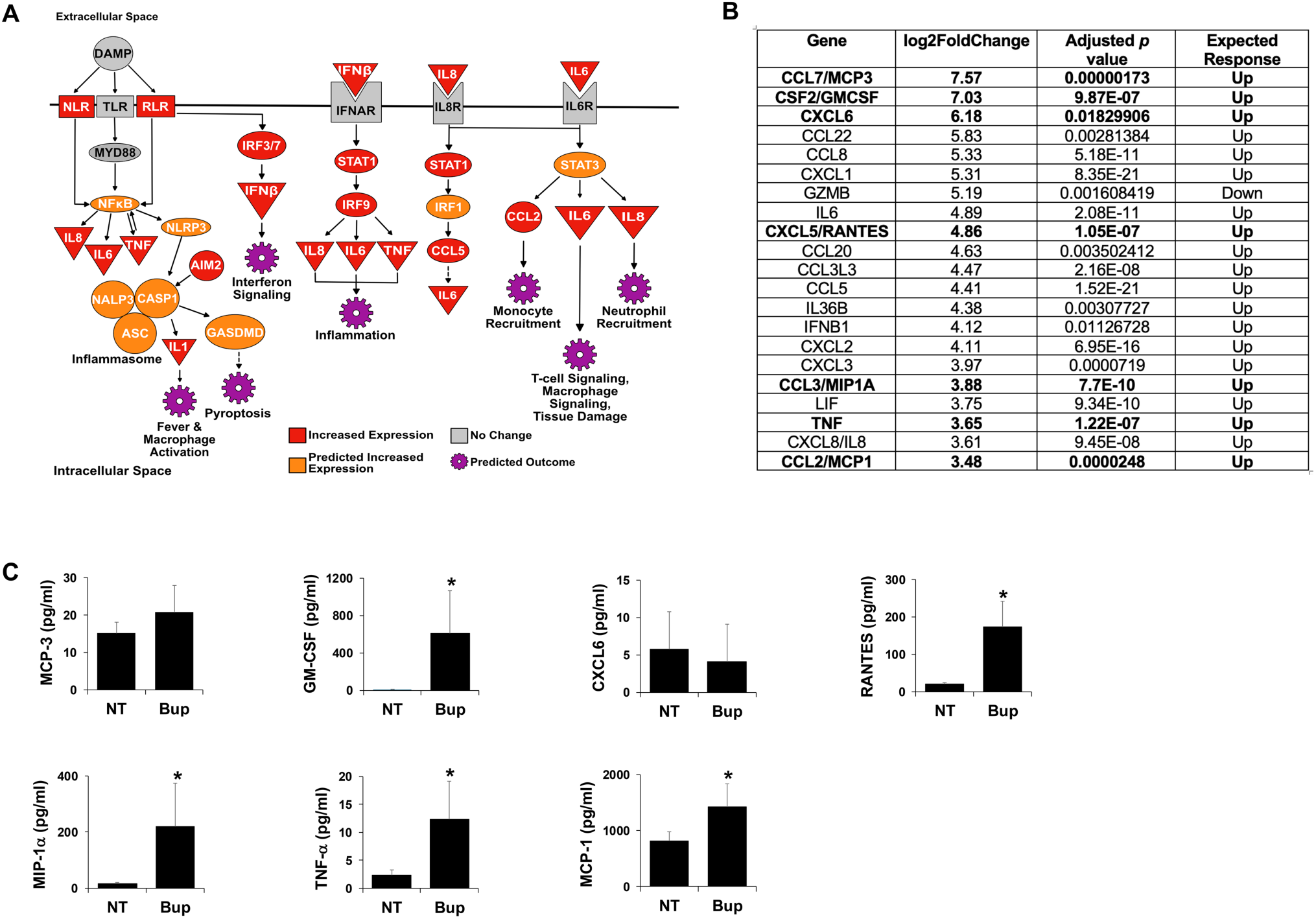
RNA-seq analysis and validation. Human FM explants from 5 patients were treated with no treatment (NT) or buprenorphine (Bup, 40μg/ml). After 48hrs total FM RNA was extracted and bulk RNA-seq analysis performed. (A) IPA analysis revealed “Cytokine Storm” in the top 5 pathways. (B) Table shows the top 20 upregulated genes from the “Cytokine Storm” IPA. The 7 bolded genes were validated by ELISA. (C) Human FM explants from 8 patients were treated with no treatment (NT) or buprenorphine (Bup, 40μg/ml). After 48hrs cell-free supernatants were collected and measured by ELISA for CCL7/MCP3, CSF2/GM-CSF, CXCL6, CXCL5/RANTES, CCL3/MIP-1α, TNF-α, and CCL2/MCP-1 (\**p*<0.05).

## Discussion

Opioid use disorder during pregnancy has increased to critical levels [1–3]. Guidelines recommend the use of methadone or buprenorphine in pregnancy because maternal and fetal outcomes are worse for untreated dyads than those who receive medications for OUD. Still, there remains a risk of preterm birth for individuals stably undergoing opiate agonist treatment. Despite the association between opioid use during pregnancy and an increased risk of preterm birth [2], no studies have attempted to understand the impact of these compounds on pregnancy outcomes, biologically or mechanistically. While there are data demonstrating a role for innate immune signaling in mediating opioid-induced inflammation in non-reproductive systems [19], this has not been explored the context of preterm birth, where there is a known role for these pathways [12, 14–16]. To address this, an established *in vitro* human explant system [13, 15, 16, 32] was used to investigate the effect of the opioid buprenorphine on FM inflammation and markers of membrane weakening, and the mechanisms involved. Herein, we report that buprenorphine triggers human FM sterile inflammation and mediators of membrane weakening through activation of TLR4/MD2, the NLRP3 inflammasome, the μ-opioid receptor, and downstream NFκB and ERK/JNK/MAPK signaling (Table 1).

**Table 1:** Summary of human FM responses to buprenorphine and the mechanisms involved. (✓) Involved; (✕) Not involved; (-) Not tested.

| | TLR4/MD2 | NLRP3 | μ-opioid<br>receptor | P65 NFκB | P38 MAPK | ERK | JNK |
| --- | --- | --- | --- | --- | --- | --- | --- |
| IL-6 | ✓ | - | ✓ | ✓ | ✓ | ✓ | ✗ |
| IL-8 | ✗ | - | ✓ | ✓ | ✗ | ✓ | ✗ |
| IL-1β | ✗ | ✓ | ✗ | ✗ | ✗ | ✗ | ✗ |
| G-CSF | ✓ | - | ✗ | ✓ | ✗ | ✓ | ✗ |
| PGE2 | ✗ | - | ✓ | ✗ | ✗ | ✗ | ✓ |
| MMP1 | ✗ | - | ✗ | ✗ | ✓ | ✓ | ✗ |

Inflammation is a major driver of preterm labor and FM inflammation is common in the setting of preterm birth. FM inflammation is known to arise from pathogenic triggers initiating inflammatory innate immune responses [14, 33, 34]. The generation of inflammatory factors can subsequently promote mediators of membrane weakening, such as PGEs and MMPs, which undermine FM integrity causing rupture, and leading to premature membrane rupture and preterm birth [35, 36]. While preterm birth is commonly associated with infection, there is growing evidence that sterile triggers can induce similar pathological inflammatory responses and pregnancy outcome [37–40], including certain medications [13]. In this study, we found that buprenorphine triggers a sterile inflammatory response in human FM explants, which mirrors the profile commonly associated with preterm birth: the inflammatory cytokine IL-6; the neutrophilic chemokine IL-8; and the inflammasome-mediated cytokine IL-1β [20, 21]. Other factors associated with membrane weakening and preterm birth were also elevated in FMs exposed to buprenorphine: PGE2, MMP1 and MMP9 [17, 18, 35, 36]. Another factor increased in FMs exposed to buprenorphine was G-CSF, which has been seen in other models of medication-induced sterile FM inflammation [13]. Our *in vitro* studies used buprenorphine at 40μg/ml which is similar to that used by others [19, 22, 23]. Recommended doses for buprenorphine during pregnancy range from 1-28mg daily, but can be higher [41, 42]. It is unclear how buprenorphine is concentrated at the maternal-fetal interface, although this medication may accumulate in the placenta [41].

Studies in non-reproductive systems demonstrated that opioids induce inflammation through innate immune signaling. Methadone increased rat glial inflammatory TNFα, MCP-1, MIP1α, and IL-1β [22] by activating TLR4 together with its adapter protein, MD2 [19, 22, 23]. Buprenorphine may also activate TLR4 and TLR2 [19, 24]. Based on this, we investigated the importance of TLR4/MD2 and TLR2 in FM responses to buprenorphine. Using specific inhibitors we found a role for TLR4/MD2 in partially mediating the FM IL-6 and G-CSF response to buprenorphine. There was no role for TLR2 in mediating the FM responses to buprenorphine. This indicated that other receptors may be involved. Indeed, we demonstrated that buprenorphine-induced FM IL-1β was mediated by the NLRP3 inflammasome, in keeping with prior studies showing an important role for this receptor complex in preterm birth [15, 43, 44]. Lastly, we tested the role of the μ-opioid receptor using the opioid receptor antagonist, (-) naloxone [28]. (-) naloxone can also inhibit TLR4/MD-2-mediated inflammation [29, 30, 45] and protects against endotoxin-induced septic shock [46]. The opioid receptor inactive isomer (+) naloxone can also inhibit TLR4/MD2-mediated inflammation [29, 30, 45] and prevents LPS and *E. coli* triggered FM inflammation and preterm birth in mice [47, 48]. In our system, opioid receptor antagonist, (-) Naloxone, partially reduced buprenorphine-induced FM secretion of IL-6, IL-8 and PGE2, indicating that both TLR4 and opioid receptor signaling are involved. It is intriguing to correlate our *in vitro* findings with clinical studies showing some improved maternal and neonatal outcomes when buprenorphine is given in combination with naloxone [49, 50]. All together, we have identified receptors involved in mediating the production of all FM inflammatory factors tested in response to buprenorphine, except for MMP1.

Since opioid receptors can activate p38 MAPK and ERK [25], and TLR4 activation triggers downstream NFκB signaling, and the MAPK pathways, p38, ERK and JNK [51], we examined these pathways in FMs exposed to buprenorphine. We found that all pathways were partially involved in regulating the FM inflammatory and membrane weakening responses to buprenorphine, although p65 NFκB and ERK signaling regulated the most factors: IL-6, IL-8, G-CSF and MMP1. p38 MAPK was involved in the FM IL-6 and MMP1 response, while JNK activation only regulated FM PGE2 production after exposure to buprenorphine.

Previous research linked FM p38 MAPK activation with premature cellular senescence and its inflammatory senescence-associated secretory phenotype as a mechanism underlying preterm premature rupture of membranes (PPROM) [31]. Since we found a role for p38 MAPK in mediating FM inflammation in response to buprenorphine, we examined senescence in our system by probing for two markers: Lamin B1 and p16. However, there was no evidence to suggest that exposing FM to buprenorphine initiated/elevated cellular senescence.

Finally, we performed RNA-seq on FM tissues to identify other FM pathways modulated by opioids and to deepen our understanding of the mechanisms underlying preterm birth associated with OUD. Our initial analysis validated our findings that buprenorphine triggered a strong FM inflammatory response; a “cytokine storm”. Inflammatory factors already identified as being upregulated were validated by our RNA-seq data. A number of other inflammatory factors were found to be elevated in buprenorphine treated FMs by RNA-seq and were subsequently validated at the protein level by ELISA. However, transcriptional increases in two factors, CCL7 and CXCL6, were not seen at the protein level, which could be due to the sensitivity of RNA-seq and low protein abundance of these factors. Regardless, most factors were validated at both the transcriptional and protein levels indicating a biologically significant inflammatory response to buprenorphine.

Strengths of this study include the use of primary human tissue explants in a highly mechanistic study. However, that this *in vitro* system models only acute exposure is also a limitation of the study. Additionally, although we demonstrated a role for TLR4/MD2, the NLRP3 inflammasome, and the μ-opioid receptor, in mediating FM inflammation in response to buprenorphine, there are likely other upstream sensors involved, and this warrants further investigation.

In summary, human FM exposure to buprenorphine results in a robust inflammatory response as well as the production of mediators of membrane weaking that together is mediated in in part by innate immune TLR4 and NLRP3 signaling, the μ-opioid receptor, and downstream NFκB and ERK/JNK/MAPK signaling. This may provide the mechanistic link between opioid use in pregnancy and the elevated risk for preterm birth, and sheds light on the experience of pregnant individuals being treated for OUD, a vulnerable and growing population. Pregnant individuals with OUD who require treatment feel faced with the difficult decision of either stopping medication and risking relapse to illness; or continuing methadone or buprenorphine therapy and increasing reproductive risks to themselves and their child. Our findings may provide a first step in being able to guide pregnant women with OUD and their providers with clinical management decision making. This study and further exploration may help identify compounds that can mitigate the inflammatory response to opioids while maintaining their ability to treat OUD.

## Author Contribution

VMA, TL, KAY and RWL designed the research. TL, MEK and EMM performed the experiments. TL, HMG and VMA analyzed the results and made the figures. TL drafted the paper. VMA, TL, KAY, RWL, and HMG revised the paper for important intellectual content.

## Acknowledgments

The authors would like to thank the Yale University Reproductive Sciences Biobank and the staff of Labor and Delivery for tissue collection.

## References

1. Forray, A., et al., Support Models for Addiction Related Treatment (SMART) for pregnant women: Study protocol of a cluster randomized trial of two treatment models for opioid use disorder in prenatal clinics. PLoS One, 2022. 17(1): p. e0261751.

2. Abdelwahab, M., et al., Risk factors for preterm birth among gravid individuals receiving buprenorphine for opioid use disorder. Am J Obstet Gynecol MFM, 2022. 4(3): p. 100582.

3. Cleveland, L.M., et al., A life-course theory exploration of opioid-related maternal mortality in the United States. Addiction, 2020. 115(11): p. 2079–2088.

4. 711, C.O.N., Opioid Use and Opioid Use Disorder in Pregnancy. Obstetrics & Gynecology, 2017. 130(2): p. e81–e94.

5. Ball, L.J., et al., Sharing of Injection Drug Preparation Equipment Is Associated With HIV Infection: A Cross-sectional Study. J Acquir Immune Defic Syndr, 2019. 81(4): p. e99–e103.

6. Committee Opinion No. 711: Opioid Use and Opioid Use Disorder in Pregnancy. Obstet Gynecol, 2017. 130(2): p. e81–e94.

7. Ecker, J., et al., Substance use disorders in pregnancy: clinical, ethical, and research imperatives of the opioid epidemic: a report of a joint workshop of the Society for Maternal-Fetal Medicine, American College of Obstetricians and Gynecologists, and American Society of Addiction Medicine. Am J Obstet Gynecol, 2019. 221(1): p. B5–B28.

8. March, o.D., 2023 Premature Birth Report Card. 2023.

9. Tita, A.T. and W.W. Andrews, Diagnosis and management of clinical chorioamnionitis. Clin Perinatol, 2010. 37(2): p. 339–54.

10. Lamont, R.F., The role of infection in preterm labour and birth. Hosp Med, 2003. 64(11): p. 644–7.

11. Goldenberg, R.L., J.C. Hauth, and W.W. Andrews, Intrauterine infection and preterm delivery. N Engl J Med, 2000. 342(20): p. 1500–7.

12. Nadeau-Vallee, M., et al., Sterile inflammation and pregnancy complications: a review. Reproduction, 2016. 152(6): p. R277–R292.

13. Fabrizio, V.A., et al., The serotonin reuptake inhibitor fluoxetine induces human fetal membrane sterile inflammation through p38 MAPK activation. J Reprod Immunol, 2023. 155: p. 103786.

14. Romero, R., et al., The role of inflammation and infection in preterm birth. Semin Reprod Med, 2007. 25(1): p. 21–39.

15. Miller, A.S., T.N. Hidalgo, and V.M. Abrahams, Human fetal membrane IL-1beta production in response to bacterial components is mediated by uric-acid induced NLRP3 inflammasome activation. J Reprod Immunol, 2021. 149: p. 103457.

16. Georges, H.M., et al., TLR8-activating miR-146a-3p is an intermediate signal contributing to fetal membrane inflammation in response to bacterial LPS. Immunology, 2024.

17. Li, W.J., et al., PGE2 vs PGF2alpha in human parturition. Placenta, 2021. 104: p. 208–219.

18. Nguyen, L.M., D.M. Aronoff, and A.J. Eastman, Matrix metalloproteinases in preterm prelabor rupture of membranes in the setting of chorioamnionitis: A scoping review. Am J Reprod Immunol, 2023. 89(1): p. e13642.

19. Gabr, M.M., et al., Interaction of Opioids with TLR4-Mechanisms and Ramifications. Cancers (Basel), 2021. 13(21).

20. Hagberg, H., C. Mallard, and B. Jacobsson, Role of cytokines in preterm labour and brain injury. BJOG, 2005. 112 **Suppl 1**: p. 16–8.

21. Christiaens, I., et al., Inflammatory processes in preterm and term parturition. J Reprod Immunol, 2008. 79(1): p. 50–7.

22. Hutchinson, M.R., et al., Possible involvement of toll-like receptor 4/myeloid differentiation factor-2 activity of opioid inactive isomers causes spinal proinflammation and related behavioral consequences. Neuroscience, 2010. 167(3): p. 880–93.

23. Hutchinson, M.R., et al., Evidence that opioids may have toll-like receptor 4 and MD-2 effects. Brain Behav Immun, 2010. 24(1): p. 83–95.

24. Jurga, A.M., et al., Blockade of Toll-Like Receptors (TLR2, TLR4) Attenuates Pain and Potentiates Buprenorphine Analgesia in a Rat Neuropathic Pain Model. Neural Plast, 2016. 2016: p. 5238730.

25. Rosenfeld, C.S., The placenta as a target of opioid drugsdagger. Biol Reprod, 2022. 106(4): p. 676–686.

26. Lim, R., Concise Review: Fetal Membranes in Regenerative Medicine: New Tricks from an Old Dog? Stem Cells Transl Med, 2017. 6(9): p. 1767–1776.

27. Lannon, S.M., et al., Synergy and interactions among biological pathways leading to preterm premature rupture of membranes. Reprod Sci, 2014. 21(10): p. 1215–27.

28. Bradberry, J.C. and M.A. Raebel, Continuous infusion of naloxone in the treatment of narcotic overdose. Drug Intell Clin Pharm, 1981. 15(12): p. 945–50.

29. Hutchinson, M.R., et al., Non-stereoselective reversal of neuropathic pain by naloxone and naltrexone: involvement of toll-like receptor 4 (TLR4). Eur J Neurosci, 2008. 28(1): p. 20–9.

30. Lewis, S.S., et al., Evidence that intrathecal morphine-3-glucuronide may cause pain enhancement via toll-like receptor 4/MD-2 and interleukin-1beta. Neuroscience, 2010. 165(2): p. 569–83.

31. Dutta, E.H., et al., Oxidative stress damage-associated molecular signaling pathways differentiate spontaneous preterm birth and preterm premature rupture of the membranes. Mol Hum Reprod, 2016. 22(2): p. 143–57.

32. Tong, M., et al., Lipopolysaccharide-Stimulated Human Fetal Membranes Induce Neutrophil Activation and Release of Vital Neutrophil Extracellular Traps. J Immunol, 2019. 203(2): p. 500–510.

33. Bakaysa, S.L., et al., Single- and double-stranded viral RNA generate distinct cytokine and antiviral responses in human fetal membranes. Mol Hum Reprod, 2014 20(7): p. 701–8.

34. Hoang, M., et al., Human fetal membranes generate distinct cytokine profiles in response to bacterial Toll-like receptor and nod-like receptor agonists. Biol Reprod, 2014. 90(2): p. 39.

35. Menon, R., et al., Diversity in cytokine response to bacteria associated with preterm birth by fetal membranes. Am J Obstet Gynecol, 2009. 201(3): p. 306 e1-6.

36. Arechavaleta-Velasco, F., et al., Production of matrix metalloproteinase-9 in lipopolysaccharide-stimulated human amnion occurs through an autocrine and paracrine proinflammatory cytokine-dependent system. Biol Reprod, 2002. 67(6): p. 1952–8.

37. Gomez-Lopez, N., et al., Intra-Amniotic Administration of HMGB1 Induces Spontaneous Preterm Labor and Birth. Am J Reprod Immunol, 2016. 75(1): p. 3–7.

38. Gomez-Lopez, N., et al., Inhibition of the NLRP3 inflammasome can prevent sterile intra-amniotic inflammation, preterm labor/birth, and adverse neonatal outcomesdagger. Biol Reprod, 2019. 100(5): p. 1306–1318.

39. Motomura, K., et al., The alarmin S100A12 causes sterile inflammation of the human chorioamniotic membranes as well as preterm birth and neonatal mortality in micedagger. Biol Reprod, 2021. 105(6): p. 1494–1509.

40. Motomura, K., et al., The alarmin interleukin-1alpha causes preterm birth through the NLRP3 inflammasome. Mol Hum Reprod, 2020. 26(9): p. 712–726.

41. Concheiro, M., et al., Maternal buprenorphine dose, placenta buprenorphine, and metabolite concentrations and neonatal outcomes. Ther Drug Monit, 2010. 32(2): p. 206–15.

42. Grande, L.A., et al., Evidence on Buprenorphine Dose Limits: A Review. J Addict Med, 2023. 17(5): p. 509–516.

43. Gomez-Lopez, N., et al., A Role for the Inflammasome in Spontaneous Preterm Labor With Acute Histologic Chorioamnionitis. Reprod Sci, 2017. 24(10): p. 1382–1401.

44. Chang, Y., et al., Association between interleukin-6 and preterm birth: a meta-analysis. Ann Med, 2023. 55(2): p. 2284384.

45. Thomas, J.H.L., et al., Toll-like receptors change morphine-induced antinociception, tolerance and dependence: Studies using male and female TLR and signalling gene KO mice. Brain Behav Immun, 2022. 102: p. 71–85.

46. Law, W.R. and J.L. Ferguson, Naloxone alters organ perfusion during endotoxin shock in conscious rats. Am J Physiol, 1988. 255(5 Pt 2): p. H1106-13.

47. Chin, P.Y., et al., Novel Toll-like receptor-4 antagonist (+)-naloxone protects mice from inflammation-induced preterm birth. Sci Rep, 2016. 6: p. 36112.

48. Chin, P.Y., et al., Toll-Like Receptor-4 Antagonist (+)-Naloxone Confers Sexually Dimorphic Protection From Inflammation-Induced Fetal Programming in Mice. Endocrinology, 2019. 160(11): p. 2646–2662.

49. Bates, M.J., et al., Household concepts of wellbeing and the contribution of palliative care in the context of advanced cancer: A Photovoice study from Blantyre, Malawi. PLoS One, 2018. 13(8): p. e0202490.

50. Petrich, M., et al., Comparison of neonatal outcomes in pregnant women undergoing medication-assisted treatment of opioid use disorder with methadone or buprenorphine/naloxone. J Matern Fetal Neonatal Med, 2022. 35(26): p. 10481–10486.

51. Guha, M. and N. Mackman, LPS induction of gene expression in human monocytes. Cell Signal, 2001. 13(2): p. 85–94.

