## Supplemental Figure 1 for "Buprenorphine Induces Human Fetal Membrane Sterile Inflammation"

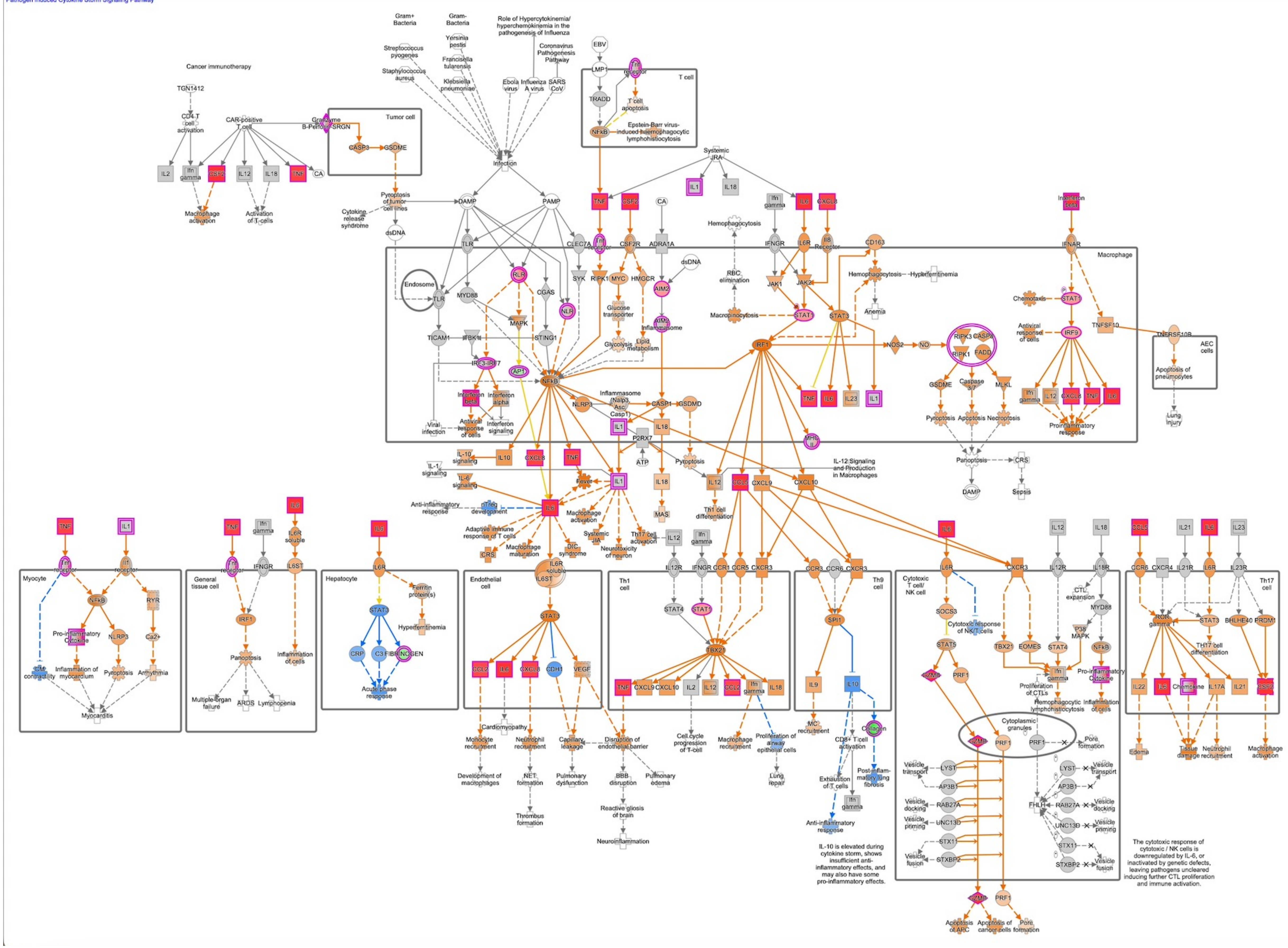


**Supplemental Figure 1. RNA-seq analysis.** Human FM explants from 5 patients were treated with no treatment (NT) or buprenorphine (Bup, 40$\mu$g/ml). After 48hrs total FM RNA was extracted and bulk RNA-seq analysis performed. IPA analysis revealed “Cytokine Storm” in the top 5 pathways.
